# The necroptosis machinery mediates axonal degeneration in a model of Parkinson disease

**DOI:** 10.1101/539700

**Authors:** Maritza Oñate, Alejandra Catenaccio, Natalia Salvadores, Cristian Saquel, Alexis Martinez, Ines Moreno-Gonzalez, Nazaret Gamez, Paulina Soto, Claudio Soto, Claudio Hetz, Felipe A. Court

## Abstract

Parkinson’s disease (PD) is the second most common neurodegenerative condition, characterized by motor impairment due to the progressive degeneration of dopaminergic neurons in the substantia nigra and depletion of dopamine release in the striatum. Accumulating evidence suggest that degeneration of axons is an early event in the disease, involving destruction programs that are independent of the survival of the cell soma. Necroptosis, a programmed cell death process, is emerging as a mediator of neuronal loss in models of neurodegenerative diseases. Here, we demonstrate activation of necroptosis in postmortem brain tissue from PD patients and in a toxin-based mouse model of the disease. Inhibition of key components of the necroptotic pathway resulted in a significant delay of 6-hydroxydopamine dependent axonal degeneration of dopaminergic and cortical neurons *in vitro*. Genetic ablation of necroptosis mediators MLKL and RIPK3, as well as pharmacological inhibition of RIPK1 *in vivo*, decreased dopaminergic neuron degeneration, improving motor performance. Together, these findings suggest that axonal degeneration in PD is mediated by the necroptosis machinery, a process here referred to as *necroaxoptosis*, a druggable pathway to target dopaminergic neuronal loss.

## Introduction

Parkinson’s disease (PD) is the second most common neurodegenerative disease, affecting 1-2% of people over 60. The increase in life span and changes in life style predicts that the burden of PD will double over the next generation (Lau and Breteler, 2006). The characteristic motor symptoms of PD include resting tremor, bradykinesia, rigidity of the limbs and abnormal gait (Goldman and Postuma, 2014). The impairment of motor control is associated to the loss of dopaminergic neurons in the substantia nigra pars compacta (SNpc) and the resulting reduction of dopamine release by these neurons in the striatum (Dauer and Przedborski, 2003). Several genetic alterations have been identified in rare familial PD cases (Corti et al., 2019), whereas more than 90% of the cases are considered idiopathic. The accumulation of abnormal protein aggregates is a transversal feature of PD, associated with the presence of Lewy bodies, protein inclusions contained misfolded forms of alpha-synuclein (Soto and Pritzkow, 2018).

A characteristic feature of PD is a progressive “dying back” process of neuronal loss. PD pathology initiates in the striatal terminals and proceeds in a retrograde fashion to the somas located in the SNpc (Kramer and Schulz-Schaeffer, 2007; Kordower et al., 2013; Orimo et al., 2008; Li et al., 2009). Studies using genetic manipulation demonstrated that apoptosis inhibition prevents cell death of mesencephalic neurons, without a significant impact in the loss of axons and dopamine depletion in the striatum (von Coelln et al., 2001; Eberhardt et al., 2000; Ries et al., 2008), suggesting that axonal degeneration occurs on a compartmentalized process in PD models. Axonal degeneration is a regulated process that occurs independent of the survival of the cell soma, executed by local mechanisms that disassemble axonal structures leading to the generation of small fragments that are then engulfed by glial cells and circulating macrophages (Salvadores et al., 2017). Among the mechanisms of axonal destruction, key components include calcium release from the endoplasmic reticulum, mitochondrial dysfunction, and ROS production (Court and Coleman, 2012). We have recently demonstrated that axonal degeneration triggered by glutamate excitotoxicity *in vitro* is commanded by a regulated program that involves components of the necroptosis machinery (Hernandez et al., 2018).

Necroptosis is a programmed cell death mechanism currently associated with cell death in several pathological conditions such as ischemia-reperfusion injury in the heart, liver injury, viral infection, cancer and neurodegeneration (Shan et al., 2003; Tonnus and Linkermann, 2017). Necroptosis can be initiated by a variety of stimuli often related to inflammation (Wallach et al., 2016), including TNF-α, Fas, TRAIL, interferons and activation of TLR by LPS, dsRNA, DNA damage, viral infection, among others (review in Grootjans et al., 2017). TNF-α-induced necroptosis is the best characterized necroptotic initiator to date. Upon TNFR1 activation by TNF-α, complex I is formed by the recruitment of the kinase RIPK1, cIAPs1/2 and adapter proteins TRADD and TRAF1/2 (Weinlich et al., 2017). In this complex, RIPK1 is poly-ubiquitylated by cIAPs (Bertrand et al., 2008) activating stress pathways (i.e. MAPK and NF-ĸB), impacting cell survival and inflammation (Kondylis et al., 2017). Nevertheless, dissociation of complex I leads to the formation of a cytosolic pro-cell death machinery (complex II) or the necrosome complex if caspases are inhibited (pharmacologically or by c-FLIP_S_) or absent (Petrie et al., 2019). Upon necrosome formation, RIPK1 is activated and autophosphorylated, activating RIPK3 which binds and phosphorylates the pseudokinase MLKL, the most downstream effector of necroptosis (Grootjans et al., 2017). Phosphorylated MLKL oligomerizes and translocates to the plasma membrane, forming pores to directly execute a necrotic form of cell death (Petrie et al., 2019; Sun et al., 2012). The mechanisms by which MLKL triggers necroptosis are still under debate, but possibly depend on the cell type affected (Gong et al., 2017; Petrie et al., 2017; Zhang et al., 2016a). Other studies suggested that MLKL interacts with the cation channel TRPM7, leading to abnormal calcium influx leading to cell swelling and plasma membrane rupture (Cai et al. 2014). Additionally, it has been demonstrated that MLKL is required for necrosome translocation to the mitochondria to enhance aerobic respiration and mitochondrial ROS production, leading to a metabolic collapse that results in non-apoptotic cell death (Yang et al., 2018).

Recent reports associated the activation of necroptosis to several neurodegenerative conditions (reviewed in Yuan et al., 2019). Inhibition of necroptosis with small molecules or genetic ablation of RIPK1, RIPK3 or MLKL exert neuroprotective effects in models of brain damage, including ischemia (Qu et al., 2016; Yin et al., 2015; Zhang et al., 2016b), traumatic injury (You et al., 2008), viral infections (Bian et al., 2017), in addition to contribute to retinal damage (Dong et al., 2012; Kim et al., 2016; Viringipurampeer et al., 2014) and spinal cord injury (Liu et al., 2015). Recent advances in the field have demonstrated the therapeutic potential of inhibiting necroptosis in several neurodegenerative diseases including amyotrophic lateral sclerosis (ALS) (Ito et al., 2016; Re et al., 2014) multiple sclerosis (MS) (Ofengeim et al., 2015) and Alzheimer’s disease (AD) (Caccamo et al., 2017). Besides, necroptosis may be relevant to control dopaminergic neuron loss in PD. Analysis of postmortem brain tissue derived from PD patients indicated increased levels of RIPK1, RIPK3 and MLKL at the SNpc (Iannielli et al., 2018). Inhibition of RIPK1 using the small molecule nec-1s protected dopaminergic neurons on a pharmacological model of PD *in vivo* (Iannielli et al., 2018) and *in vitro* (Wu et al., 2015). Nevertheless, whether necroptosis activation is functionally associated to motor dysfunction and denervation in PD has not been explored.

In this study, we investigated the contribution of the necroptotic pathway to axonal degeneration in dopaminergic neurons in the context of PD. Analysis of human PD postmortem brain tissue and mouse models of the disease indicated the activation of key components of the necroptosis machinery in dopaminergic neurons of the SNpc. Functional assessment indicated that targeting RIPK1 or MLKL significantly reduced axonal degeneration in dopaminergic, as well as cortical primary neurons. Genetic ablation of RIPK3 and MLKL attenuated axonal degeneration, translating into an improvement of motor performance. To assess the therapeutic potential of necroptosis to PD, we administrated nec-1s to a mouse model of PD and observed clear neuroprotection at the histological and behavioral level. Our results demonstrate a novel function of the necroptosis machinery in controlling the mechanisms of axonal destruction in PD and suggest that strategies to inhibit necroptosis may have important therapeutic benefits to attenuate neurodegeneration in PD.

## Results

### 1 Pharmacological inhibition of the necroptosis machinery delays 6-OHDA-induced neurite degeneration *in vitro*

We first studied the involvement of necroptosis in PD using primary neuronal cultures from the embryonic mesencephalon, as this region give rise to the SNpc during brain development. 6-OHDA was used as a relevant neurotoxic PD insult. We inhibited RIPK1 using nec-1s, a strategy that prevents the formation of the necrosome complex (Takahashi et al., 2012). Mesencephalic neuronal cultures were exposed to 6-OHDA or vehicle as control, in the presence or absence of nec-1s followed by morphological assessment by immunostaining. 6-OHDA induced the degeneration of neurites characterized by fragmentation of the cytoskeleton and neurite beading, a phenomenon that was completely prevented by nec-1s treatment (Fig. 1A). Quantitative assessment of degeneration (See Supplementary Fig. 1A and method section) confirmed the protective effects achieved by nec-1s over neurites (Fig. 1B). Further classification of neurites into intact, beaded or fragmented indicated almost complete protection of their integrity (Fig. 1C). Of note, nec-1s alone did not alter neurite morphology of mesencephalic neurons (Fig. 1A-C).

**Figure 1.**
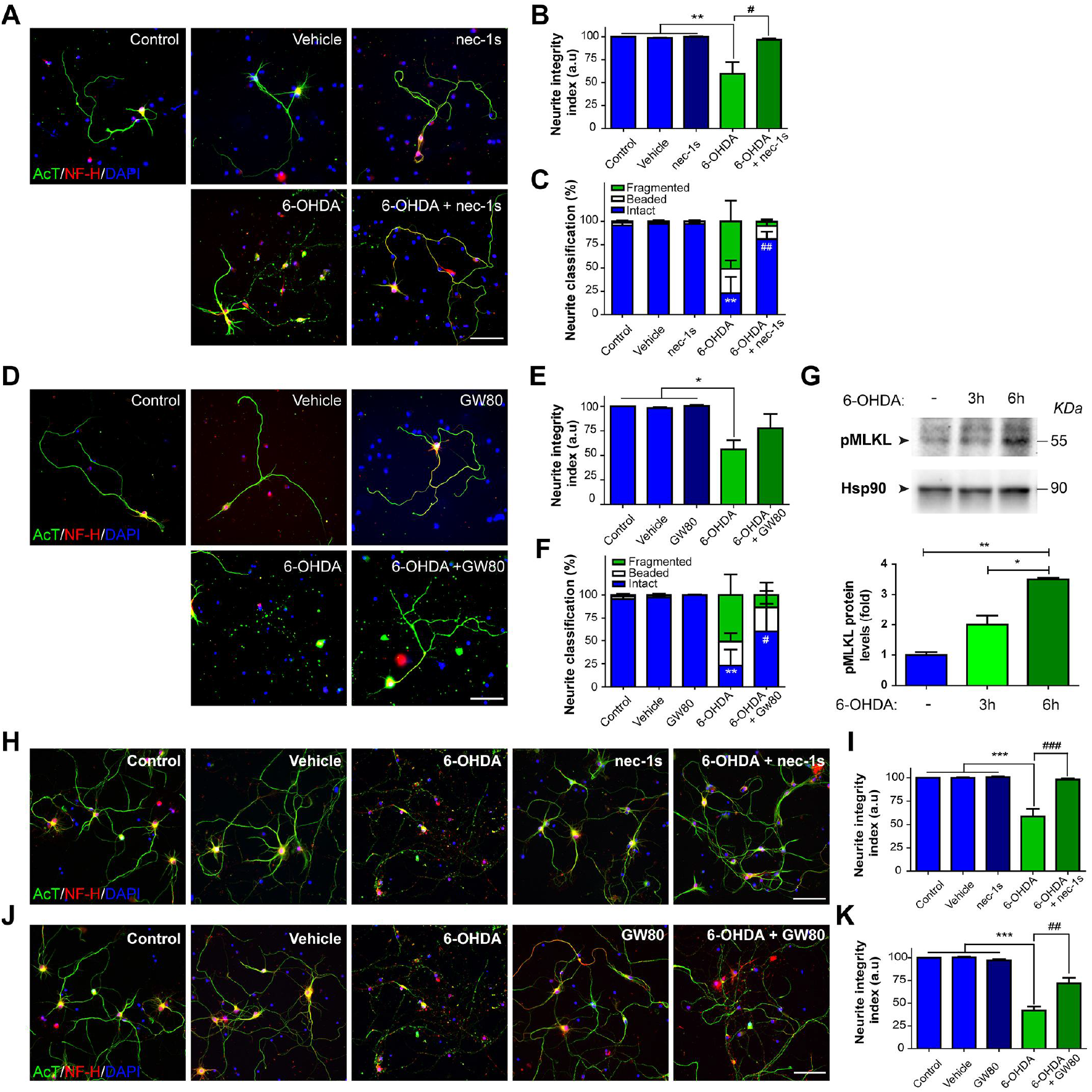
Pharmacological inhibition of RIPK1 and MLKL delay neurite degeneration *in vitro*. Mesencephalic neuronal cultures were treated for 6 hours with 6-OHDA alone (40 μM for 6 hours) or together with the RIPK1 inhibitor nec-1s (30 μM) or vehicle (**A-C**). Similar cultures were treated with 6-OHDA alone or together with the MLKL inhibitor GW80 or vehicle (**D-F**). Untreated cultures were used as control. Cells were immunostained for acetylated tubulin (AcT, green) and neurofilament heavy chain (NF-H, red). Nuclei were stained using DAPI (blue). In each condition, the neurite integrity index (**B and E**) or a classification of neurite morphology (**C and F**) was calculated. (**G**) Cortical primary cultures were treated with 6-OHDA for 3 or 6 hours. pMLKL expression was measured by western blot. Hsp90 was used as loading control. Densitometric analysis was performed in each condition for pMLKL and normalized against Hsp90. Cortical neuronal cultures were treated for 6 hours with 6-OHDA alone or together with vehicle or the RIPK1 inhibitor nec-1s (**H-I**). Similar cultures were treated with 6-OHDA alone or together with vehicle or the MLKL inhibitor GW80 (**J-K**). Cells were stained for acetylated tubulin (green), neurofilament heavy chain (NF-H) and DAPI (blue). (**I, K**) Neurite integrity index calculated for each condition. Scale bar, 50 μm. Data are shown as mean ± SEM. Statistical differences were obtained using one-way ANOVA followed by Bonferroni’s *post hoc* test. * *p* < 0.05, ** *p* < 0.01; *** *p* < 0.001 compared to control, vehicle and nec-1s or GW80 conditions. # *p* < 0.05; ## *p* < 0.01, ### *p* < 0.001 compared to 6-OHDA condition. n = 3 per group.

Since RIPK1 can trigger apoptotic cell death under certain conditions (Declercq et al., 2009), we studied the participation of MLKL, the canonical necroptotic executor protein, in 6-OHDA-dependent neurodegeneration. To this end, pharmacological inhibition of MLKL was evaluated in mesencephalic neuronal cultures using GW806742x (GW80), a compound that binds to MLKL blocking its translocation to the plasma membrane (Hildebrand et al., 2014). Although GW80 treatment induce a slight, but not significant protection in 6-OHDA treated neurons (Fig. 1D-E and Supplementary Fig. 1B), neurite classification analysis revealed a significant protection of GW80 over 6-OHDA-dependent neurite degeneration process (Fig. 1F).

In addition to dopaminergic neurons, PD has been also associated to neurodegeneration of olfactory, cortical and autonomic peripheral neurons in different stages of the disease (Braak et al., 2003). Therefore, we performed pharmacological inhibition of RIPK1 and MLKL in cortical neuronal cultures exposed to 6-OHDA. Neurons were treated with 6-OHDA for 3 and 6 hours and phosphorylation of MLKL (pMLKL) was evaluated by western blot. Low basal expression of pMLKL was detected in control conditions after vehicle treatment, which were increased by 3-fold after 6 hours of 6-OHDA treatment (Fig. 1G). Morphologically, treatment with nec-1s or GW80 resulted in a significant protection of neurite degeneration of cortical neurons exposed to 6-OHDA (Fig. 1H-K). Taken together, our results indicate that pharmacological inhibition of two key components of the necroptotic pathway reduces neurodegeneration in mesencephalic and cortical neurons cultures.

### 2 Activation of necroptosis markers in the brain of PD patients

We next assessed the possible activation of necroptosis markers in the brain of PD patients. We analyzed the phosphorylation levels of MLKL, a measure of its activation, in postmortem samples derived from PD patients and age-matched healthy controls. Analysis of MLKL phosphorylation at Ser358 and neuromelanin pigment positive cells (marker of dopaminergic neurons) in the SNpc indicated extensive MLKL phosphorylation in PD cases (Fig. 2A, see inset and arrows in PD samples). Quantitative analysis of the pMLKL staining revealed a significantly higher immunoreactivity in brain tissue derived from PD patients compared to healthy control subjects (Fig. 2B). As expected, control tissue presented marked neuromelanin staining, which was considerably lower in samples from PD patients (Fig. 2A), suggesting loss of dopaminergic neurons.

**Figure 2.**
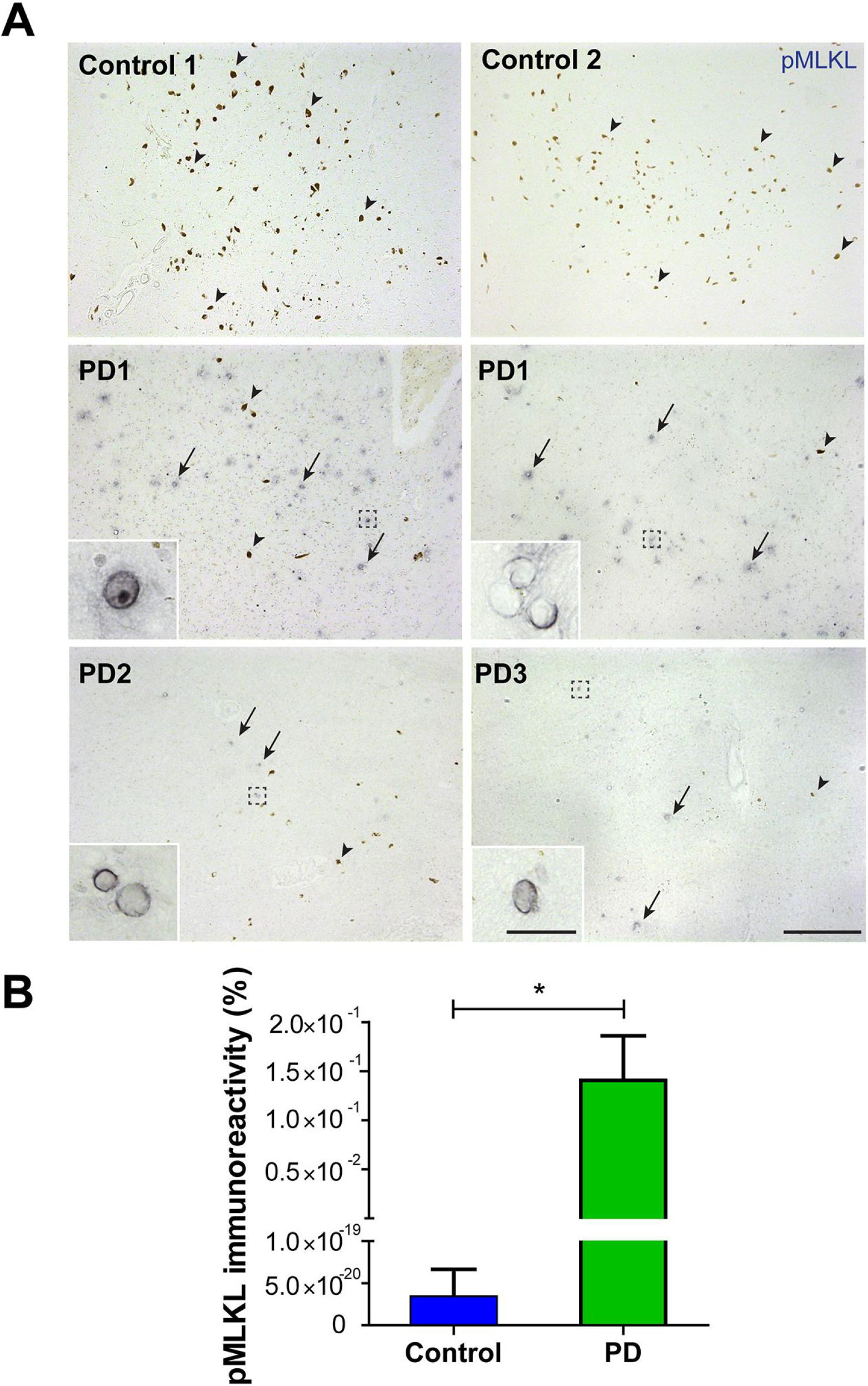
Activation of pMLKL in postmortem samples of human PD brains. (**A**) Representative images of substantia nigra from healthy control patients (HC) and Parkinson’s disease patients (PD) immunostained for pMLKL (dark blue). Neuromelanin pigmentation was used to identify dopaminergic neurons, which is seen as a brown coloration and indicated with arrowheads. pMLKL-positive neurons are indicated using arrows. (**B**) Percentage of pMLKL immunoreactivity was measured in each condition (see methods). Scale bar, 300 μm, inset, 25 μm. Data are shown as mean ± SEM. Statistical differences were obtained using Mann Whitney test. * *p* < 0.05. n = 3 per group.

### 3 MLKL activation and necrosome formation in the nigrostriatal pathway after exposure to 6-OHDA

To establish whether the necroptotic pathway contributes to dopaminergic neuron degeneration in experimental PD, we first analyzed the levels of activation of critical molecular mediators in animals injected with 6-OHDA. To this end, animals were exposed to 6-OHDA in the striatum and then biochemical and histological analysis was performed in different regions of the nigrostriatal circuit. Western blot analysis indicated elevated levels of phosphorylated MLKL in the striatum of animals only after 3 days of 6-OHDA injection and not at 7 days post-treatment (Fig. 3A). Moreover, a transient upregulation of total MLKL levels was observed at 3 days post-6-OHDA injection (Fig. 3A). Similar results were observed for pMLKL in the nigrostriatal pathway (Fig. 3B). Unexpectedly, when the SNpc of the same animals was analyzed, no changes in pMLKL and total MLKL was observed after the 6-OHDA challenge (Fig. 3C).

**Figure 3.**
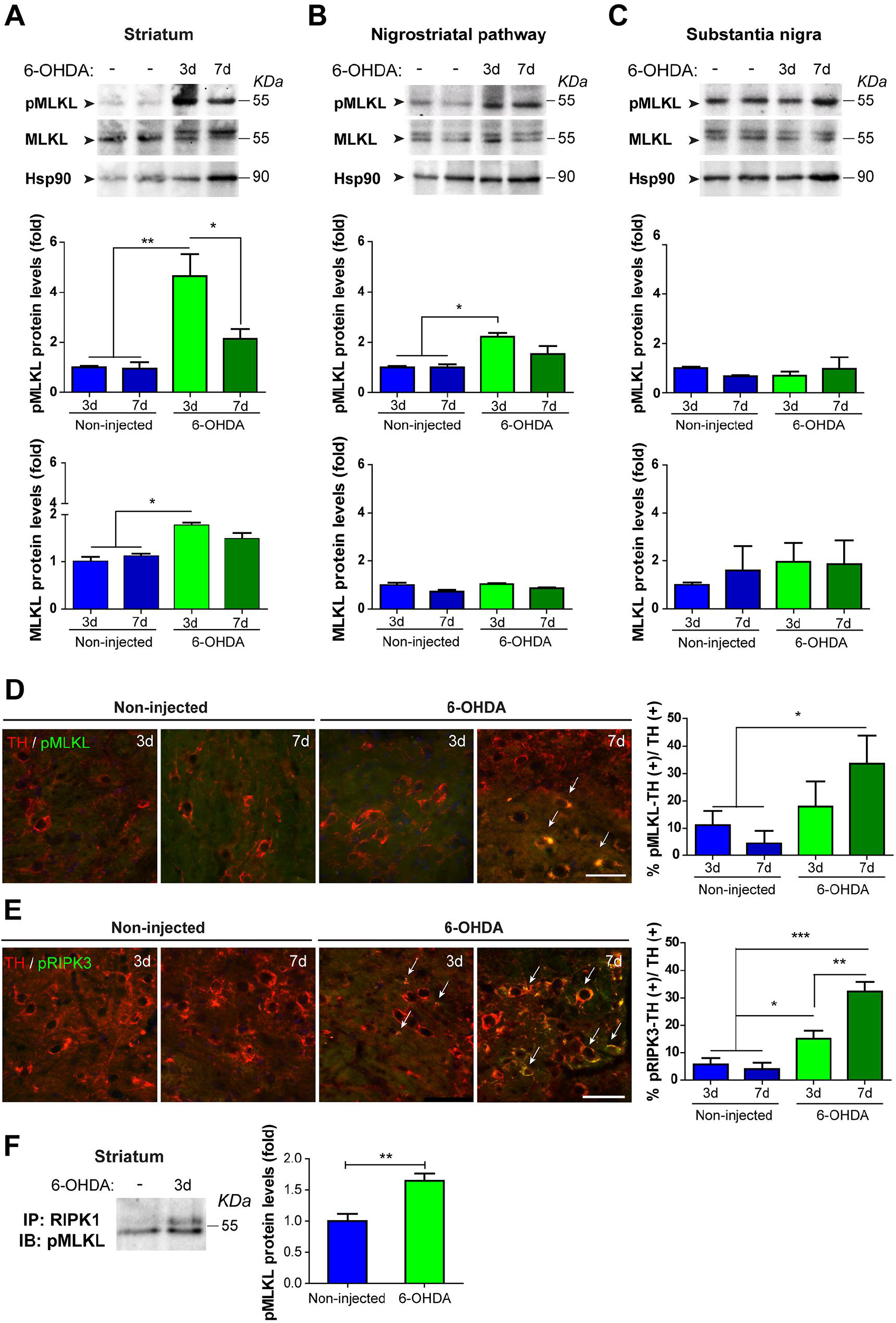
Activation of necroptosis in the nigrostriatal pathway after 6-OHDA treatment *in vivo*. Wild-type mice were injected with 6-OHDA in the right striatum and analyzed 3 or 7 days post-injection. Coronal sections of 2 mm thickness were obtained from the striatum, axonal tract and substantia nigra, and the non-injected and injected regions were divided for western blot analysis. pMLKL and MLKL protein expression was evaluated in striatum (**A**), nigrostriatal pathway (**B**) and substantia nigra (**C**). Hsp90 was used as loading control. (**D-E**) Coronal sections of substantia nigra from mice injected with 6-OHDA and studied at 3 and 7 days post injection by immunofluorescence. (**D**) Sections were immunostained for TH (red) and pMLKL (green) or (**E**) TH (red) and pRIPK3 (green). TH-positive neurons also immunoreactive for pMLKL or pRIPK3 were counted and normalized to the total dopaminergic TH+ neurons in each condition (arrows). (**F**) Proteins extracted from 3 days-injected striatum and contralateral hemisphere were immunoprecipitated with an antibody against RIPK1 and probed for pMLKL. Relative pMLKL levels were calculated by densitometry. Data are shown as mean ± SEM. Statistical differences were obtained using one-way ANOVA followed by Bonferroni’s *post hoc* test in (A), (B), (C), (D) and (E) and by student’s t-test in (F). * *p* < 0.05; ** *p* < 0.01, *** *p* < 0.001. n = 3 animals per condition.

We then determined if the activation of MLKL occurs on a cell-autonomous manner in dopaminergic neurons. Analyses of the distribution of pMLKL together with tyrosine hydroxylase (TH) using co-immunofluorescence revealed a significant increase in pMLKL in dopaminergic neurons at the SNpc (Fig. 3D). Similar results were obtained when phosphorylated RIPK3 were analyzed in the same experimental setting (Fig. 3E). Activation of MLKL by RIPK3 is dependent on the formation of a RIPK1-RIPK3-MLKL necrosome complex (Zhang et al., 2016a). Therefore, formation of the necrosome was evaluated in the striatum 3 days after 6-OHDA injection by immunoprecipitation. Pull down of RIPK1 revealed an increase in pMLKL-RIPK1 interaction in the 6-OHDA injected hemisphere compared to the contralateral side (Fig. 3F). Together, these results demonstrate a progressive and retrograde activation of the necroptosis machinery in a relevant experimental model of PD.

### 4 The necroptosis machinery contributes to axonal degeneration and neuronal loss on an animal model of PD

To study the possible participation of the necroptosis signaling pathway to axonal degeneration *in vivo*, we set up the experimental conditions to dissociate the process of early axonal degeneration from the loss of somas at the SNpc on a toxicological model of PD. We established a methodology to evaluate the nigrostriatal circuit in the mouse brain since most of the studies in PD are focused only on striatal denervation of axonal terminals in the striatum (CPu) and neuronal cell loss in the SNpc. Serial coronal sections of the entire nigrostriatal circuit of animals unilaterally injected with 6-OHDA at the CPu were obtained at 3 and 7 days post-surgery and immunostained for TH. Striatal denervation was calculated by measuring the optical density of TH in CPu sections (Fig. 4A). Axonal degeneration was analyzed along rostro-caudal axis by evaluating axonal tract length in non-injected and injected hemispheres (Fig. 4B, left). In the non-injected hemisphere, axonal tract lengths showed no differences along sections, however a progressive decrease was detected in injected hemispheres 3 and 7 days after 6-OHDA injection (Supplementary Fig. 2A-B). Estimation of the percentage of axonal loss in each section at 3 and 7 days post 6-OHDA injection demonstrated a spatial and temporal progression of axonal tract degeneration (Fig. 4B, right). Finally, the loss of dopaminergic neuronal somas was estimated in the SNpc by quantification of TH-positive neurons (Fig. 4C, left). No changes were observed across the SNpc at 3 days post 6-OHDA injection, whereas 7 days of treatment resulted in significant neuronal loss (Fig. 4C, right). These results were also validated when the spatial distribution of TH neurons was quantified (Supplementary Fig. 2C). Together, these results showed that the 6-OHDA model is suitable to study axonal degeneration in the absence of evident neuronal loss.

**Figure 4.**
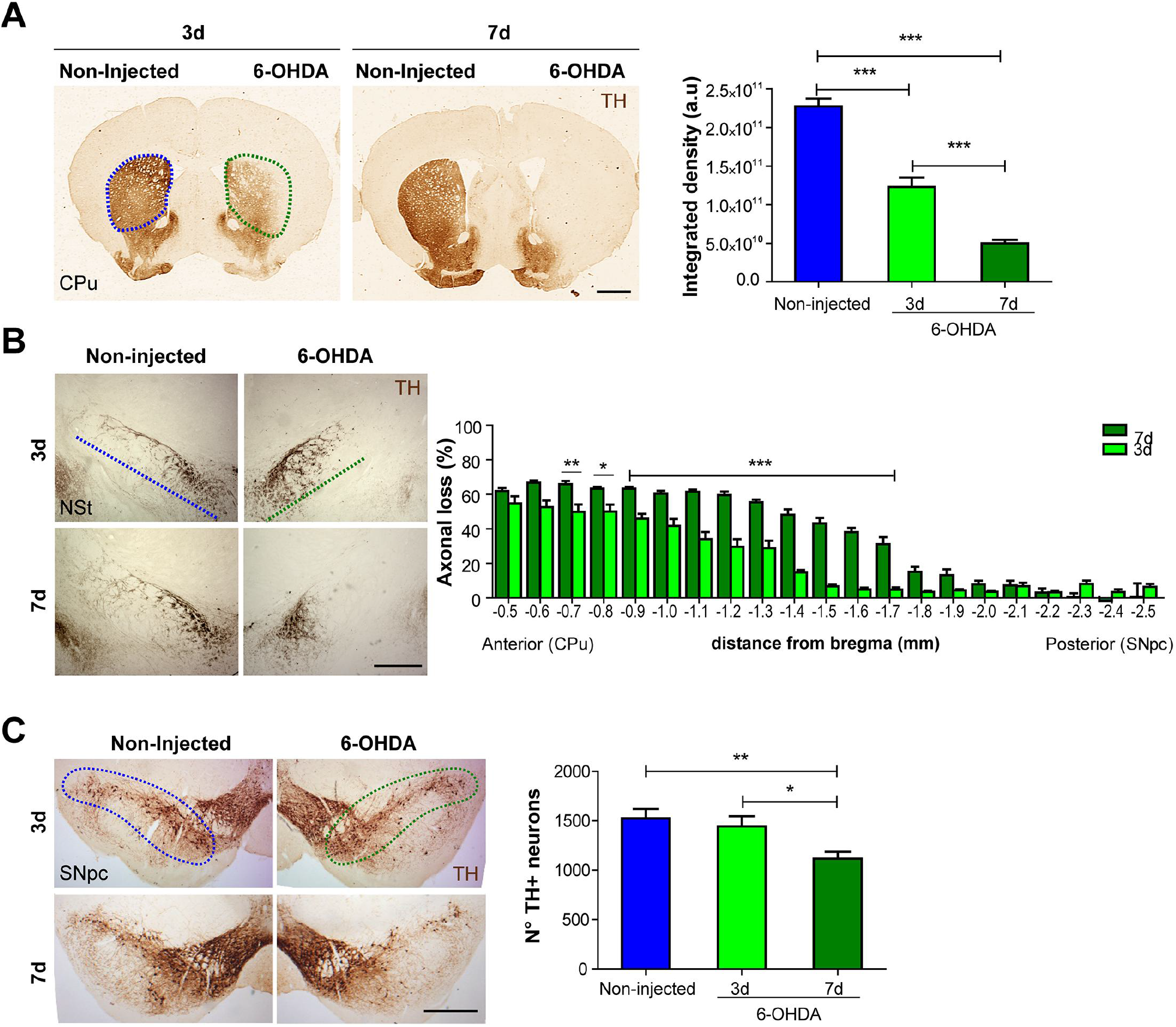
6-OHDA treatment induces a progressive and retrograde degeneration of nigrostriatal neurons. Wild-type mice were injected with 6-OHDA in the right striatum. The contralateral hemisphere was kept non-injected as a control. Serial coronal sections of the entire nigrostriatal circuit were obtained at 3 and 7 days post 6-OHDA injection and immunostained for TH. The non-injected and injected striatum from each hemisphere are demarcated with blue and green dashed lines, respectively. (**A, left**) Striatal area analyzed (CPu) at 3 and 7 days after injection. Scale bar, 1 mm. (**A, right**) Striatal denervation was calculated as the total integrated optical density in each condition. (**B, left**) Nigrostriatal (NSt) pathway at 3 and 7 days post-injection. Axonal tracts are indicated using dashed lines for each hemisphere at 3 and 7 days post 6-OHDA injection. Scale bar, 500 μm. (**B, right**) Spatial distribution of axonal loss at 3 and 7 days post 6-OHDA injection. (**C, left**) Images from the substantia nigra pars compacta (SNpc) at 3 and 7 days post 6-OHDA injection. Scale bar, 500 μm. (**C, right**) Quantification of the total number of TH-positive cells in the entire SNpc at 3 and 7 days post 6-OHDA injection. Data are shown as mean ± SEM. Statistical differences were obtained using one-way ANOVA in (A) and (C) and twoway ANOVA in (B) followed by Bonferroni’s *post hoc* test. * *p* < 0.05; ** *p* < 0.01; *** *p* < 0.001. n = 10 animals per group.

We next performed loss-of-function studies to define the involvement the necroptosis machinery in axonal neurodegeneration and motor impairment triggered by 6-OHDA. To this end, we genetically disrupted the expression of MLKL, as the participation of this protein defines a canonical necroptotic process. MLKL-deficient mice (MLKL^−/−^) and their littermate control animals (MLKL^+/+^) were injected with 6-OHDA in the right striatum and after 7 days. Optical density analysis of the striatum showed no differences in striatal denervation in MLKL^−/−^ animals when compared to control mice (Fig. 5A). In sharp contrast, ablation of MLKL expression significantly protected dopaminergic axonal tracks in animals challenged with 6-OHDA (Fig. 5B and Supplementary Fig. 3A-B). At the SNpc, the number of TH-positive somas were protected in MLKL^−/−^ mice against 6-OHDA, observing a 21% of loss when compared to a reduction of 34% in littermate control animals (Fig. 5C and Supplementary Fig. 3C). To validate these results with a second key necroptotic player, we genetically targeted the expression of RIPK3. RIPK3-deficient mice (RIPK3^−/−^) and their littermate control animals (RIPK3+/+) were injected with 6-OHDA in the right striatum and 7 days after, serial coronal sections of the entire circuit were obtained to evaluate axonal degeneration. Again, optical density analysis of the striatum showed no differences in striatal denervation in RIPK3^−/−^ animals when compared to control mice (Fig. 5D). In sharp contrast, ablation of RIP3K expression significantly protected dopaminergic axonal tracks in animals challenged with 6-OHDA (Fig. 5E and Supplementary Fig. 3D-E). Together, these results suggest that MLKL and RIPK3 contributes to the dying back degeneration of axons observed in the 6-OHDA model.

**Figure 5.**
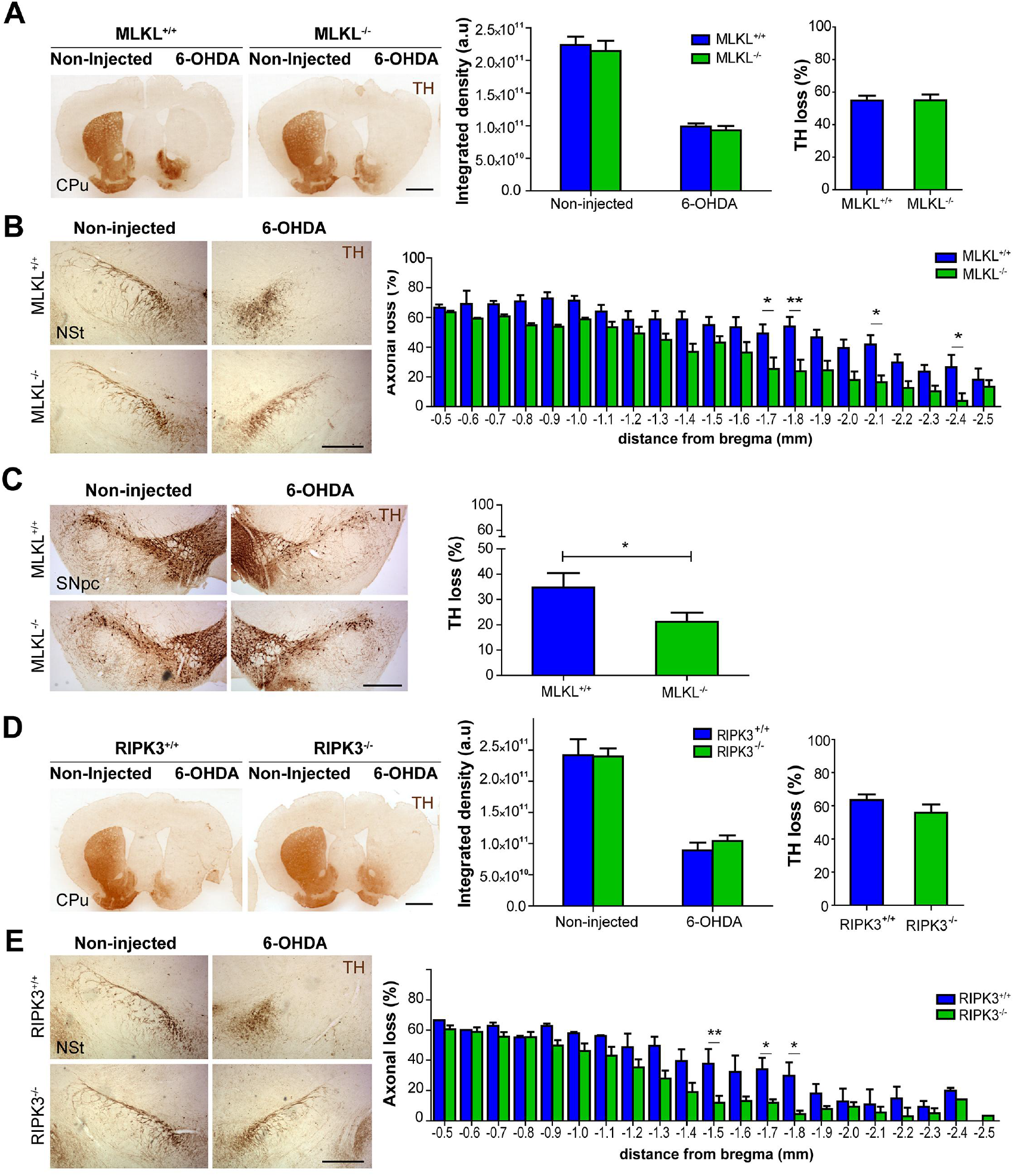
MLKL and RIPK3 deficiency delays neurodegeneration and motor impairment after 6-OHDA injection. MLKL^−/−^, RIPK3^−/−^ and corresponding WT sibling mice were injected with 6-OHDA in the right striatum (CPu) and the contralateral hemisphere was kept non-injected as a control. Serial coronal sections of the entire nigrostriatal circuit were obtained 7 days after 6-OHDA injection and immunostained for TH. (**A, left**) Representative striatal coronal sections from MLKL^−/−^ and MLKL^+/+^ mice unilaterally injected with 6-OHDA in the right striatum. Scale bar, 1 mm. (**A, right**) Striatal denervation was calculated as total integrated optical density in non-injected and injected hemisphere. Percentage of TH loss staining was estimated from integrated density. (**B, left**) Representative images from nigrostriatal axons (NSt) from MLKL^−/−^ and MLKL^+/+^ mice injected and immunostained for TH. Scale bar, 500 μm. (**B, right**) Spatial distribution of axonal loss in each genotype. (**C, left**) 6-OHDA induced neuronal loss in MLKL^−/−^ and MLKL^+/+^ mice was analyzed in the substantia nigra pars compacta (SNpc). Scale bar, 500 μm. (**C, right**) Quantification of total number of TH-positive cells in the entire SNpc represented as the percentage of neuronal loss in each genotype. (**D, left**) Representative striatal coronal sections from RIPK3^−/−^ and RIPK3^+/+^ mice injected with 6-OHDA in the right striatum. Scale bar, 1 mm. (**D, right**) Striatal Integrated density and percentage of TH loss calculated in each condition. (E, left) Representative coronal sections from nigrostriatal pathway in RIPK3^−/−^ and RIPK3^+/+^ mice injected. (**E, right**) Spatial distribution of axonal loss in each genotype. Data are shown as mean ± SEM. Statistical differences were analyzed using two-way ANOVA followed by Bonferroni’s *post hoc* test in (A and D, for Integrated density measurements, B and E, right), and by student’s t-test in (A and D, for percentage of loss). * *p* < 0.05; ** *p* < 0.01. n = 8 animals per group.

### 5 Ablation of MLKL and RIPK3 expression improve motor performance on a PD model

Given that both MLKL and RIPK3 deficiency reduced axonal neurodegeneration induced by 6-OHDA *in vivo*, we determined if these effects translated in the recovery of the motor capacity. The cylinder test was performed to measure forepaw akinesia after unilateral 6-OHDA lesion. No differences were found in RIPK3^−/−^ however, MLKL^−/−^ mice showed a significant improvement in forepaw akinesia at 7 days after injection (Fig. 6A-B). To monitor motor coordination, we performed the rotarod test in the same animals before surgery, and at 3 and 7 days after 6-OHDA injection. No basal alterations in motor performance were detected in both MLKL and RIPK3 null mice (Fig. 6C-D). Remarkably, at 3 days post injection, a reduced decay in performance of about 40% was observed in MLKL^−/−^ mice compared to control animals (Fig. 6C). Similarly, improved motor control was observed in RIPK3 knockout (Fig. 6D). Together, these results indicate that the necroptosis machinery mediates in part the neurodegeneration cascade observed in our toxicological model of PD, resulting in improved motor activity.

**Figure 6.**
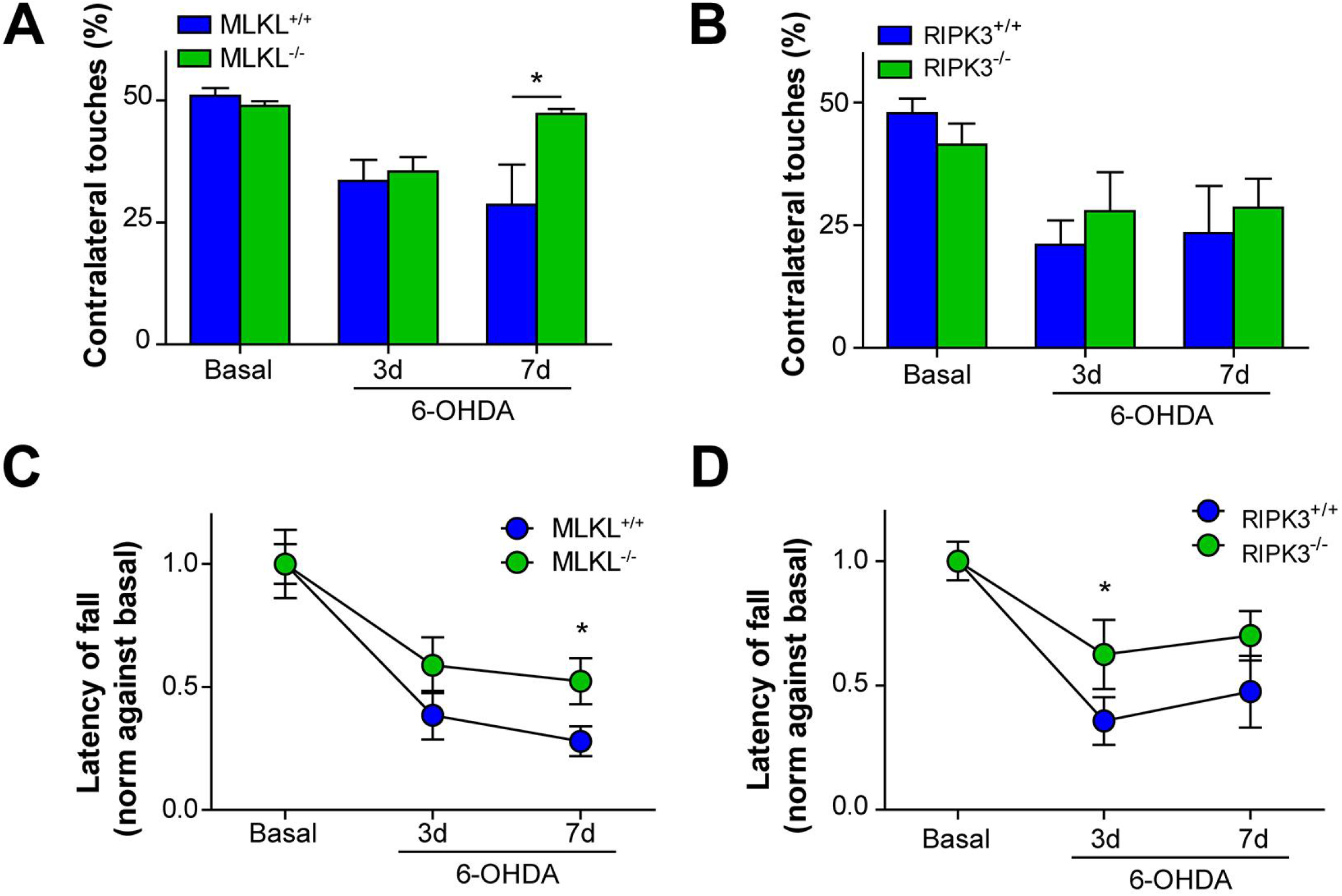
MLKL and RIPK3 ablation improves motor behavior after 6-OHDA injection. MLKL^−/−^, RIPK3^−/−^ and WT littermate mice were injected with 6-OHDA in the right striatum. Contralateral hemisphere was kept non-injected as a control. (**A, B**) Forepaw akinesia after injection was performed using the cylinder test. Percentage of touches from the paw contralateral to the injection side was measured in MLKL^−/−^ and RIPK3^−/−^, respectively. (**C, D**) Motor performance was tested using the rotarod test by measuring the latency to fall in an accelerated protocol in each genotype. Data are shown as mean ± SEM. Statistical differences were analyzed using two-way repeated measures ANOVA followed by Bonferroni’spost *hoc* test. * *p* < 0.05. n = 8 animals per group.

### 6 Pharmacological inhibition of RIPK1 decreases dopaminergic neuron degeneration and reduces motor impairment of experimental PD

Our previous results are indicative of a functional role of the necroptosis machinery in the degeneration of dopaminergic axons in *in vivo*. To increase the translational potential of these findings, we tested the consequences of pharmacologically inhibiting the necroptosis machinery in experimental PD. To this end, we intraperitoneally administrated nec-1s daily for three days before and after exposing animals to the 6-OHDA challenge at the striatum. 7 days after 6-OHDA injection morphological analysis of the brain was performed. Consistent with our genetic studies, treatment of animals with nec-1s have no effect in 6-OHDA-dependent striatal denervation (Fig. 7A). When the nigrostriatal axonal tract was visualized in the same animals, a significant protection was observed after nec-1s administration (Fig. 7B and Supplementary Fig. 4A-B). Furthermore, a significant decrease in forepaw akinesia was found in nec-1s treated mice at 3 days post 6-OHDA injection (Fig. 7C). Finally, motor performance was improved in animals injected with nec-1s (Fig. 7D). Together, these results indicate that pharmacological intervention of the necroptosis machinery results in significant neuroprotection of experimental PD.

**Figure 7.**
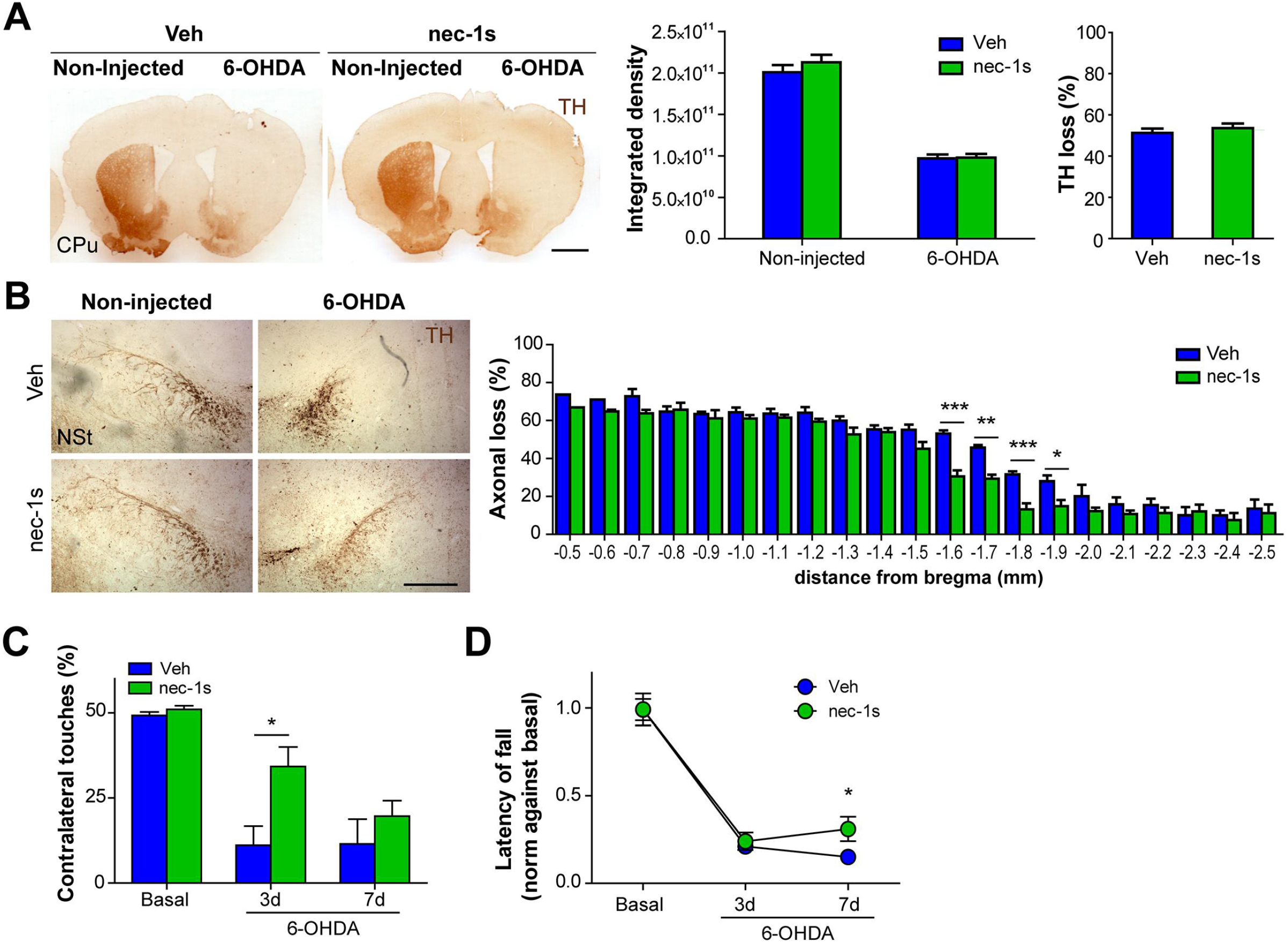
Pharmacological inhibition of RIPK1 contributes to neurodegeneration and motor impairment after 6-OHDA injection. WT mice were pre-treated with nec-1s for 3 days and the unilaterally injected with 6-OHDA in the right striatum. Then mice were followed by 7 days with daily injections of nec-1s. Vehicle (Veh) treatment was used as a control for the nec-1s injections. Left striatum was kept un-injected as control. Serial coronal sections of the entire nigrostriatal circuit were obtained 7 days after 6-OHDA injection and immunostained for TH. (**A, left**) Representative striatal (CPu) coronal sections from Veh or nec-1s treated mice unilaterally injected with 6-OHDA. Scale bar, 1 mm. (**A, right**) Striatal denervation 7 days after 6-OHDA injection was calculated as the total integrated optical density in non-injected and injected hemispheres from Veh or nec-1s treated mice. The percentage of TH loss staining was estimated from integrated density. (**B, left**) Representative images from nigrostriatal axons (NSt) from WT mice unilaterally injected with 6-OHDA and treated with Veh or nec-1s. Scale bar, 500 μm. (**B, right**) Spatial distribution of axonal loss 7 days after 6-OHDA and treated with Veh or nec-1s. (**C**) Forepaw akinesia was evaluated using the cylinder test. Percentage of touches from the paw contralateral to the injection side was measured. (**D**) Motor performance was tested using the rotarod test by measuring the latency to fall in an accelerated protocol. Data are shown as mean ± SEM. Statistical differences were analyzed using two-way ANOVA followed by Bonferroni’s *post hoc* test in (A for Integrated density), by student’s t-test in (A for percentage of TH loss), and by two-way repeated measures ANOVA followed by Bonferroni’s *post hoc* test in (B, C and D). * *p* < 0.05. n = 9 animals per group.

## Discussion

PD is a chronic neurodegenerative condition characterized by the degeneration of nigrostriatal dopaminergic neurons located in the SNpc. Axonal loss is emerging as a critical pathological event in PD, which occurs in early stages of the disease and precedes somatic neuronal death (Cheng et al., 2010; Chung et al., 2009; Kordower et al., 2013; Tagliaferro and Burke, 2016; Tagliaferro et al., 2015). In addition, axonal degeneration is a common feature of other diseases including AD, ALS, and MS, representing an interesting transversal target for therapeutic interventions. However, the exact mechanisms by which axons degenerate in PD remains unknown. Here, we unveiled a functional role for RIPK1, RIPK3 and MLKL as part of the molecular machinery that executes axonal degeneration in dopaminergic neurons upon an oxidative stress injury. Since the axonal compartment corresponds to a cellular extension, we propose that necroptosis *per se* is not a mechanism of axonal destruction, rather the necroptotic machinery has an alternative function in disassembling cellular structures. Motoneurons and dopaminergic neurons have the largest volume occupied by dendrites and axons, where the volume of the soma is virtually a minor portion of the whole cell (Matsuda et al., 2009). Because of the compartmentalized nature of neurons, the idea that the necroptosis machinery controls the “death” of the axon may have an evolutionary origin where components of necrosome further specialized in this alternative disassembling pathway. Based in our findings, we would like to propose the concept of “necroaxoptosis” as a novel mechanism of axonal degeneration mediated by components of the necroptosis machinery.

Axonal loss in models of PD has been usually quantified using relatively simple measurements or restricted to delimited sampling areas, not including the entire nigrostriatal tract (Cheng et al., 2010 West, 2013). Here, we developed a novel histological method to study of the whole nigrostriatal pathway along the antero-posterior axis. Remarkably, measurements of axonal length in every section was sufficient to depict the retrograde and progressive effect of the 6-OHDA injection along the tract at the morphological level. In addition, biochemical analysis demonstrated a coincident activation of the necroptosis pathway along the temporal and spatial dimensions.

Activation of necroptosis has been described in several neurodegenerative conditions (Shao et al., 2017; Zhang et al., 2017). Importantly, interaction between RIPK1, RIPK3 and MLKL is detected in postmortem samples from MS and AD patients. In our toxicological PD model, we observed an early increase in the interaction between RIPK1 and pMLKL in the striatum, suggesting the formation of the necrosome complex. By using pMLKL as a proxy for necroptosis activation, we have found a progressive activation of the pathway after 6-OHDA damage from terminals in the striatum to axonal tracts in the nigrostriatal pathway and finally somas in the SNpc, which coincides with data obtained in human postmortem samples from PD patients. Whether necroaxoptosis contributes to axonal degeneration in other diseases where necroptosis is important to induce the death of the cell soma (i.e. AD, ALS, brain ischemia and MS) remains to be determined.

Previous evidence in neurodegenerative conditions indicates that necroptosis can be activated in glial cells, including astrocytes and oligodendrocytes (Ito et al., 2016; Ofengeim et al., 2015; Re et al., 2014). Our *in vitro* data suggested that activation of necroptosis takes place in neurons after treatment with 6-OHDA, which is also in agreement with a cell-autonomous activation of necroptosis in the axonal compartment initiated by different pro-degenerative stimuli including excitotoxicity and inhibition of microtubule-dependent transport (Hernandez et al., 2018). Multiple pathological processes may activate necroptosis in PD, including inflammation and oxidative stress. Its known that α-synuclein released from degenerating neurons stimulates microglia, leading to microglia activation and TNF-α secretion (Collins et al., 2012 Croisier et al., 2005). In agreement with this idea, MPTP-induced striatal dysfunction is reduced in mice lacking TNF-α expression, whereas the loss of dopaminergic neurons at the SNpc is not affected (Ferger et al., 2004). In other neurodegenerative conditions, including MS and ALS, sustained inflammation has been associated to TNFα-mediated necroptosis (Wegner et al., 2017; Yuan et al., 2019). In the experimental autoimmune encephalomyelitis (EAE) mouse model of MS, oligodendrocytes were found to be highly sensitive to TNFα-induced necroptosis, whereas in microglia, TNFα induces inflammation as they express high levels of RIPK1 and low levels of MLKL (Ofengeim et al., 2015). Similarly, in a mouse model of ALS, oligodendrocytes showed greater sensitivity to TNFα-induced cell death, and microglia showed increased phosphorylation of RIPK1 and increased expression of a number of inflammatory genes (Ito et al., 2016). Therefore, in diverse neurodegenerative conditions, activation of RIPK1 plays a major role in microglia-dependent inflammatory activation, leading to a cell non-autonomous induction of necroptosis in oligodendrocytes and as a consequence, neurodegeneration (Wegner et al., 2017). Whereas the occurrence of necroptosis in oligodendrocytes across the axon surface contributes to necroaxoptosis remains to be determined.

Together, evidence suggest that the necroptotic machinery operates on a cell-autonomous manner in neurons to execute axonal degeneration possibly involving ROS production as a triggering event (Hernandez et al., 2018), a pathological event involved in PD (Hsu et al., 2000; Ko et al., 2000; Orth et al., 2004; Surmeier et al., 2011). Additionally, a cell non-autonomous mechanism may contribute to this degeneration process in dopaminergic neurons where TNFα might be secreted by microglia. Since the mechanisms associated to axonal loss are different from those involved in canonical programmed cell death and the destruction of the cell body, necroaxoptosis emerges as a novel and potential mechanism for axonal degeneration. Thus, inhibition of necroaxoptosis appears to be a promising therapeutic target for functional and structural preservation of axons and terminals in PD and in other neurodegenerative conditions.

## Materials and Methods

### Neuronal primary cultures

Mesencephalic neuronal cultures were obtained from embryonic E14.5 C57Bl/6 mice and cortical neuronal cultures were obtained from embryonic E18.5 Sprague-Dawley rats. The protocol used for both type of primary culture was the same, except the brain structure dissected. Briefly, meninges were removed from each brain and ventral mesencephalon or cortex from both hemispheres were dissected, trypsinized and plated onto 0.1 mg/mL poly-L-lysine coverslips or plastic dishes. Neurons were grown in Neurobasal media supplemented with B27 and L-glutamine. After 7 days *in vitro* (DIV), neuronal cultures were treated with 40 μM of 6-OHDA (in 0.2% ascorbic acid) or vehicle. For inhibitory treatments, cells were exposed to 30 μM of necrostatin-1s (nec-1s, Biovision) or 0.5 μM of GW806742x (GW80, AdipoGen). For control conditions, cells were incubated with fresh Neurobasal supplemented medium (control), vehicle or inhibitors alone.

### Neurite integrity index

Quantification of neurite integrity in *in vitro* experiments was calculated as the ratio between staining area of intact neurites and total staining area (area of intact neurites + area of fragmented neurites) of Acetylated Tubulin immunofluorecense. Non-neuronal staining was discarded by co-localization of Acetylated Tubulin and Neurofilament heavy chain staining. All images were acquired and processed simultaneously using Image J software. Binary masks of each image were obtained to analyze size fragment of particles. Particles with a size area equal or lower than 25 μm^2^ and with a circularity index higher than 0.3 were classified as degenerated neurite fragments. Particles with a size area higher than 25 μm^2^ with a circularity index lower than 0.3 were classified as intact neurite.

### Human tissue staining

Human brain tissue (substantia nigra pars compacta) from PD patients (n=3) and control samples (n=3) was obtained from Banner Sun Health Research Institute. After blocking the endogenous peroxidase activity with 3% H_2_O_2_-10% methanol for 20 min, brain sections were heated at 80°C for 30 min in 50mM citrate buffer pH 6.0 for antigen retrieval prior overnight incubation with anti-phospho-MLKL (S358) mouse monoclonal antibody (1:200: Signalway antibody). Primary antibody was detected by incubating 1 hour with sheep anti-mouse HRP-linked secondary antibody (General Electric), and peroxidase reaction was visualized using DAB Kit (Vector) following the manufacturer’s instructions. Finally, all sections were dehydrated in graded ethanol, cleared in xylene, and cover-slipped with DPX mounting medium. 3 samples from each individual were examined under a bright field microscope (DMI6000B, Leica Microsystems) and representative photomicrographs were taken with a digital camera (DFC310 FX Leica). Immunoreactivity percentage was defined as the percentage of area stained with anti-pMLKL related to the substantia nigra analyzed in different coronal sections (3 sections/subject). pMLKL-immunopositive signal was converted into 8-bit gray scale and identified by a threshold intensity to quantify the area labeled per total area analyzed. Human samples were manipulated following the universal precautions for working with human specimens and as directed by the Institutional Review Board of the University of Texas Health Science Center at Houston (HSC-MS-14-0608).

### Experimental animals

Adult (12-16 weeks old) C57BL/6 mice, MLKL knockout and RIPK3 knockout mice were used. MLKL and RIPK3 knockout mice were kindly provided by Dr. Douglass Green (St. Jude Children’s Research Hospital, Memphis, TN, USA) and have been described previously (Nogusa et al., 2016; Wu et al., 2013). Animals were kept under standard conditions of light and temperature and were feed with food and water *ad libitum* in the Animal Facility of the Sciences Faculty of the Mayor University. The research protocol n° 08-2016 was approved by the Animal Care and Use Scientific Ethic Committee of the Mayor University.

### Toxicological model of PD

Mice were anesthetized with isoflurane and placed in a stereotaxic frame (David Kopf Instruments, USA). A single unilateral injection was performed in the right striatum at the following coordinates: anteroposterior (AP): +0.07 cm, medio-lateral (ML): −0.17 cm and dorso-ventral (DV): −0.31 cm, relative to bregma (according to Mouse Brain Atlas, Paxinos and Franklin, 2008) as previously described (Castillo et al., 2015). 2 μl of a solution of 8 μg of 6-OHDA (4 μg/μl in 0.2% ascorbic acid) was injected at a rate of 0.5 μl/min. Animals were euthanized by overdose of anesthesia at different days post injection for further analysis.

### Histological analysis

Mice were deeply anesthetized with isoflurane and intracardially perfused with isotonic saline followed by 4% paraformaldehyde. Brains were dissected, post-fixed overnight in 4% paraformaldehyde at 4 °C and then incubated in 30% sucrose. Tissue was cryoprotected in optimal cutting temperature compound (OCT, Tissue-Tek) at −20 °C and serial coronal sections of 25 μm thick containing the nigrostriatal circuit (from rostral striatum to ventral midbrain) were obtained using a cryostat (Leica, CM1860). Injected hemisphere was marked for identification. Serial free-floating sections were processed for immunohistochemistry as previously described (Castillo et al., 2015). Briefly, slices were quenched with 0.3% H_2_O_2_ for 30 min, blocked with 0.5% bovine serum albumin and 0.2% triton X-100 for 2 hours and incubated with primary antibody (rabbit anti-Tyrosine hydroxylase, Millipore) overnight at 4 °C. Then, sections were washed with 0.1 M PBS and incubated with secondary biotinylated antibody (goat anti-rabbit, Vector Laboratories) for 2 hours at RT. After washing, slices were incubated with avidin-biotin-peroxidase complex (Vector Laboratories) for 1 hour at RT followed by 0.1 M PBS washes and developed with 3,3-diaminobenzidine (DAB, Sigma-Aldrich). Finally, sections were mounted on glass slides with Entellan medium (Merck).

### Densitometry analysis of dopaminergic striatal innervation

Dopaminergic terminals at the striatum (CPu) were analyzed every 100 μm of the entire area. Sections were scanned using an Epson Perfection V600 Photo scanner and striatal innervation was quantified by measuring the optical density of TH-immunoreactivity in the striatum using ImageJ software (NIH, USA). Results were expressed as the integrated density of the entire region and as the percentage of TH immunoreactivity loss compared to control hemisphere as previously described (Castillo et al., 2015).

### Quantification of dopaminergic axons

Dopamine axonal tract at the nigrostriatal pathway (NSt) images were obtained in a Nikon Eclipse E200 microscope. The length of the TH-positive axonal tract was calculated in each sections every 100 μm. Spatial distribution of axonal length was calculated in each section along the antero-posterior axis in injected and contralateral non-injected hemisphere. Percentage of axonal loss was calculated in the injected hemisphere compared with the contralateral hemisphere in each section. Measurements were performed using Image J software.

### Dopaminergic neuron cell counting

Estimation of the number of TH-positive neurons at the SNpc was performed in serial sections of the entire midbrain every 100 μm as previously reported (Castillo et al., 2015). The percentage of TH-immunoreactive neurons relative to the contralateral (non-injected) side was determined by counting the total number of TH-positive neurons in the entire SNpc. Spatial distribution analysis of each section was performed along antero-posterior axis. Measurements were performed using Image J software.

### Tissue processing and co-immunoprecipitation

For biochemical analysis, brains were extracted and washed in ice-cold 0.1 M PBS and then sectioned in a stainless steel brain matrix (coronal slices, 1 mm spacing). 2 mm sections from striatum, nigrostriatal axonal pathway and mesencephalon (containing the entire SNpc) were homogenized in 100 μl of RIPA buffer (50 mM Tris-HCl, 150 mM NaCl, 1% NP-40, 0.5% Sodium deoxycholate, 0.1% SDS and 5 mM EDTA, pH 7.6) containing protease (1 mM PMSF and protein inhibitor cocktail, PIC) and phosphatase inhibitors (1 mM NaF and 50 mM Na_3_VO_4_). Protein concentration was estimated using the Pierce BCA Protein Assay Kit (Thermo Scientific). For cell culture experiments, cells were collected in 0.1 M PBS and then homogenated in RIPA buffer for protein quantification as previously described. For immunoprecipitation experiments, 100 μg of brain protein lysates were incubated with 4 μg of anti-RIPK1 (Cell signaling) for 48 hours while rotating at 4 °C. After incubation, 50 μl of Protein A magnetic beads (Invitrogen) were added to each sample and were incubated at 4 °C for 3 h while rotating. Following magnetic separation, beads were mixed with loading buffer and boiled at 90 °C for 5 min. Samples were loaded onto a 10% SDS/PAGE gel and Western blot was performed for pMLKL antibody as followed described.

### Western blot

Brain and cell culture lysates containing 50 μg of protein were loaded into 10% SDS/PAGE gels and transferred onto methanol-activated PVDF membranes (Thermo Scientific). Membranes with transferred proteins were blocked/permeabilized for 1 hour in 5% BSA in TBS and incubated with different antibodies (mouse anti-pMLKL, Millipore, rabbit anti-MLKL, Abcam, and rabbit anti-Hsp90, Santa Cruz) overnight at 4 °C. Membranes were incubated with HRP-secondary antibodies for 1 hour at RT and revealed using ECL (Invitrogen) and Chemidoc™ MP Imaging System (Biorad). For densitometry analysis of the bands, Image J software was used.

### Culture cells immunofluorescence

For immunocytochemical analysis, neurons were fixed with 4% paraformaldehyde for 15 min. Then, cells were blocked-permeabilized in 5% gelatin from cold water fish skin and 0.1% triton X-100 for 1 h followed by incubation with primary antibodies (mouse anti-acetylated tubulin, and rabbit anti-Neurofilament heavy chain, Sigma-Aldrich) overnight at 4 °C. After washing with 0.1 M PBS, cells were incubated with secondary antibodies (goat anti-mouse Alexa Fluor 488 and goat anti-rabbit Alexa Fluor 546, Thermo-Fisher) for 2 hours at RT, then washed with 0.1 M PBS and coverslips were mounted on slides with Fluoromont-G (Electron Microscopy Sciences) solution with DAPI nuclear staining (Thermo-Fisher). Images were obtained using a Leica DMi8 microscope.

### Mouse brain immunofluorescence

For immunofluorescence analysis, 25 μm coronal sections from the substantia nigra pars compacta were used. Antigen retrieval was performed with boiling Sodium Citrate 10mM, pH 6 for 10 min and then, sections were blocked/permeabilized in 5% BSA, 0,3% Triton X-100 in 0.1M TBS for 2 hours at RT, following by primary antibody incubation overnight at 4°C (rabbit anti-pRIPK3, Abcam; mouse anti-TH, Millipore; mouse anti-pMLKL: Millipore rabbit anti-TH, Millipore in blocking/permeabilizing solution). Sections were washed in 0.1M TBS and then incubated in secondary antibodies for 2 hours at RT in TBS (goat anti-rabbit Alexa Fluor 488, goat anti-mouse Alexa Fluor 546, goat anti-mouse Alexa Fluor 488 and goat anti-rabbit Alexa Fluor 546, Thermo-Fisher). Finally, sections were washed in 0.1M TBS and mounted in Mowiol (Sigma 81381).

### In vivo necrostatin-1s treatment

For nec-1s (Biovision, CA, USA) preparation the compound was dissolved in DMSO (50% w/v) and then transferred into 35% PEG solution as previously described (Ofengeim et al., 2016). C57Bl/6 mice were treated for 3 days with nec-1s (8 mg/Kg i.p.) or vehicle before surgery and then, daily injected for 7 days after 6-OHDA injection.

### Behavioral tests in mice

Behavioral tests were performed in injected animals before surgery (baseline) and at 3 and 7 days post-injection for analysis of functional motor and coordination performance. The cylinder test was used to assess asymmetric forelimb use as previously reported (Castillo et al, 2015). Mice were placed in a transparent glass cylinder of 20 cm diameter for 5 min and was videotaped during the test. The number of ‘ipsilateral’ (right touches) and ‘contralateral touches’ (left touches) was quantified and represented as the percentage of ‘contralateral touches’ of all movements observed. For the rotarod test, mice were placed into a rotating rod (Model LE8500, Panlab SL) and the time until mice fell was measured (latency fall). Animals were subjected to 4 trials per day using the accelerated speed test protocol, consisted in increasing speed trial starting with 4 rpm up to 40 rpm within 120 s. Animals waited ~5 min between each trial to avoid fatigue.

### Statistical analysis

Data are shown as mean ± SEM. Statistical analysis were performed using Student’s t-test or one-way ANOVA, two-way ANOVA test or two-way repeated measures ANOVA test, followed by Bonferroni’s *post hoc* test or with Mann Whitney non-parametric test, using GraphPad Prism 5.0 software.

## Acknowledgments

We are grateful to the Banner Sun Health Research Institute Brain and Body Donation Program of Sun City, Arizona for the provision of brain tissue. This work was supported by Geroscience Center for Brain Health and Metabolism FONDAP-15150012 (FAC and CH), Ring Initiative ACT1109 (FAC and CH), FONDECYT-1150766, Canada-Israel Health Research initiative, jointly Funded by the Canadian Institutes of Health Research, the Israel Science Foundation, the international Development Research Centre, Canada and the Azrieli Foundation, Canada (FC), Conicyt Doctoral Fellowship 21130843 (MO), FONDECYT 1140549 (CH), Millennium Institute P09-015-F (CH), European Commission R&D MSCA-RISE 734749 (CH). Michael J Fox Foundation for Parkinson’s Research – Target Validation grant 9277 (CH), FONDEF ID16I10223 (CH), FONDEF D11E1007 (CH), US Office of Naval Research-Global N62909-16-1-2003 (CH), U.S. Air Force Office of Scientific Research FA9550-16-1-0384 (CH), ALSRP Therapeutic Idea Award AL150111 (CH), Muscular Dystrophy Association 382453 (CH), and CONICYT-Brazil 441921/2016-7 (CH).

## Authors Contributions

M.O., A.C., N.S., C.S., I.M-G., N.G., P.S., performed experiments M.O, A.C., C.S, A.M, N.G. analyzed data. M.O., I.M-G., C.S., C.H., F.A.C., designed experiments and wrote the manuscript.

## Conflict of Interest

The authors declare no conflict of interest.

