## Supplementary Figures for "The necroptosis machinery mediates axonal degeneration in a model of Parkinson disease"

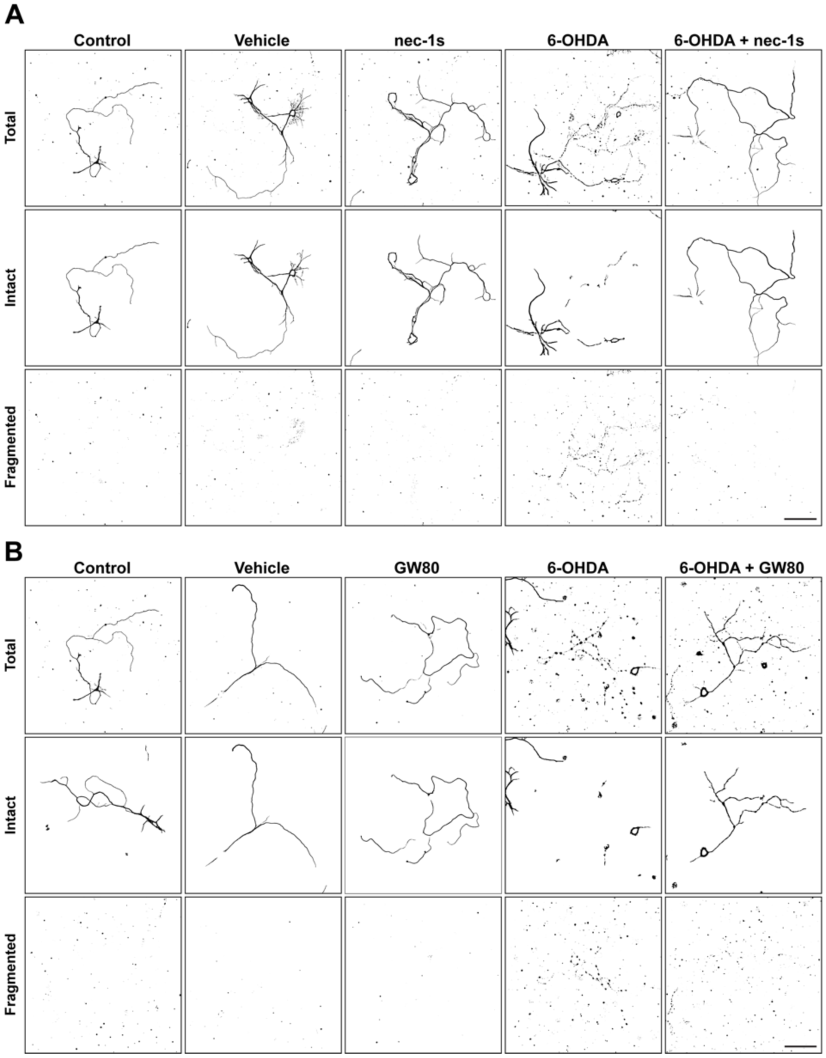


**Supplementary Figure 1. Effect of necroptosis inhibition *in vitro* after 6-OHDA treatment (related to Figure 1). (A)** Representative Acetylated tubulin (AcT) binary masks from images of mesencephalic neuronal cultures were treated for 6 hours with 6-OHDA alone or together with vehicle or the RIPK1 inhibitor nec-1s. **(B)** Binary masks from images of mesencephalic cultures treated for 6 hours with 6-OHDA alone or together with vehicle or the MLKL inhibitor GW80. Using imageJ and the particle analyzer macro, setting parameters for bigger and continuous fragments were used to obtain intact binary masks (middle panels) and setting parameters for smaller and fragmented particles were used to obtain fragmented binary masks (lower panel). Scale bars, 100 µm.

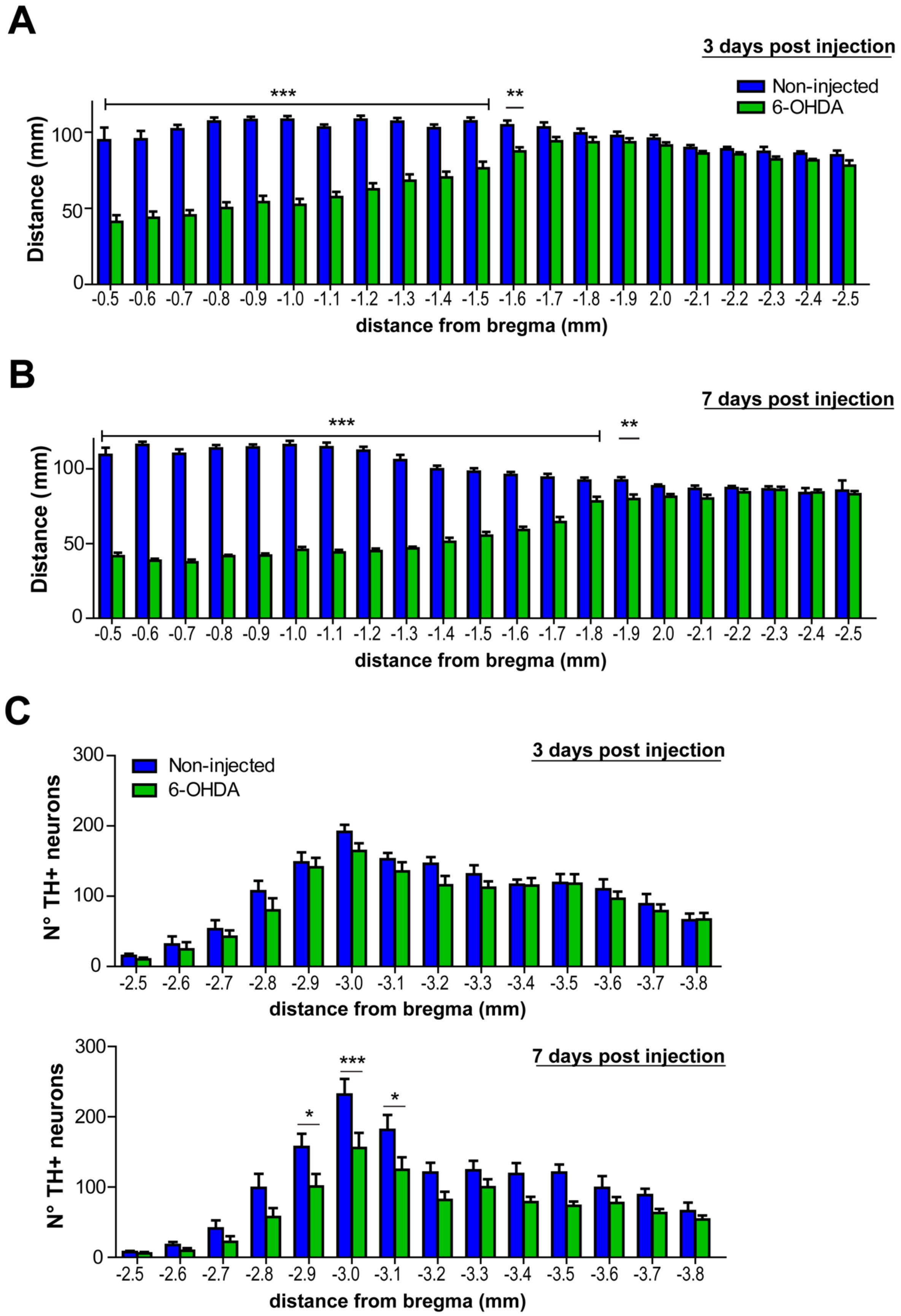
A
3 days post injection
Non-injected
6-OHDA
***
**
Distance (mm)
0
50
100
-0.5
-0.6
-0.7
-0.8
-0.9
-1.0
-1.1
-1.2
-1.3
-1.4
-1.5
-1.6
-1.7
-1.8
-1.9
2.0
-2.1
-2.2
-2.3
-2.4
-2.5
distance from bregma (mm)
B
7 days post injection
***
**
Distance (mm)
0
50
100
-0.5
-0.6
-0.7
-0.8
-0.9
-1.0
-1.1
-1.2
-1.3
-1.4
-1.5
-1.6
-1.7
-1.8
-1.9
2.0
-2.1
-2.2
-2.3
-2.4
-2.5
distance from bregma (mm)
C
3 days post injection
Non-injected
6-OHDA
N° TH+ neurons
0
100
200
300
-2.5
-2.6
-2.7
-2.8
-2.9
-3.0
-3.1
-3.2
-3.3
-3.4
-3.5
-3.6
-3.7
-3.8
distance from bregma (mm)
7 days post injection
*
***
*
N° TH+ neurons
0
100
200
300
-2.5
-2.6
-2.7
-2.8
-2.9
-3.0
-3.1
-3.2
-3.3
-3.4
-3.5
-3.6
-3.7
-3.8
distance from bregma (mm)

**Supplementary Figure 2. Analysis of axonal loss after 6-OHDA injection (related to Figure 4).** WT mice were unilateral injected with 6-OHDA in the right striatum. Left hemisphere was used as non-injected control side. Axonal lengths were analyzed in serial coronal sections immunostained for TH every 100 µm at 3 **(A)** and 7 **(B)** days. Antero-posterior coordinates relative from bregma are shown in the Y axis. **(C)** Spatial distribution of the number of TH-positive neurons in the SNpc in the non-injected and injected hemisphere at 3 (upper graph) and 7 (lower graph) days post 6-OHDA injection. Data are shown as mean ± SEM. Statistical differences were analyzed using two-way ANOVA followed by Bonferroni's *post hoc* test. * $p < 0.05$, ** $p < 0.01$; *** $p < 0.001$. n = 10 animals per group.

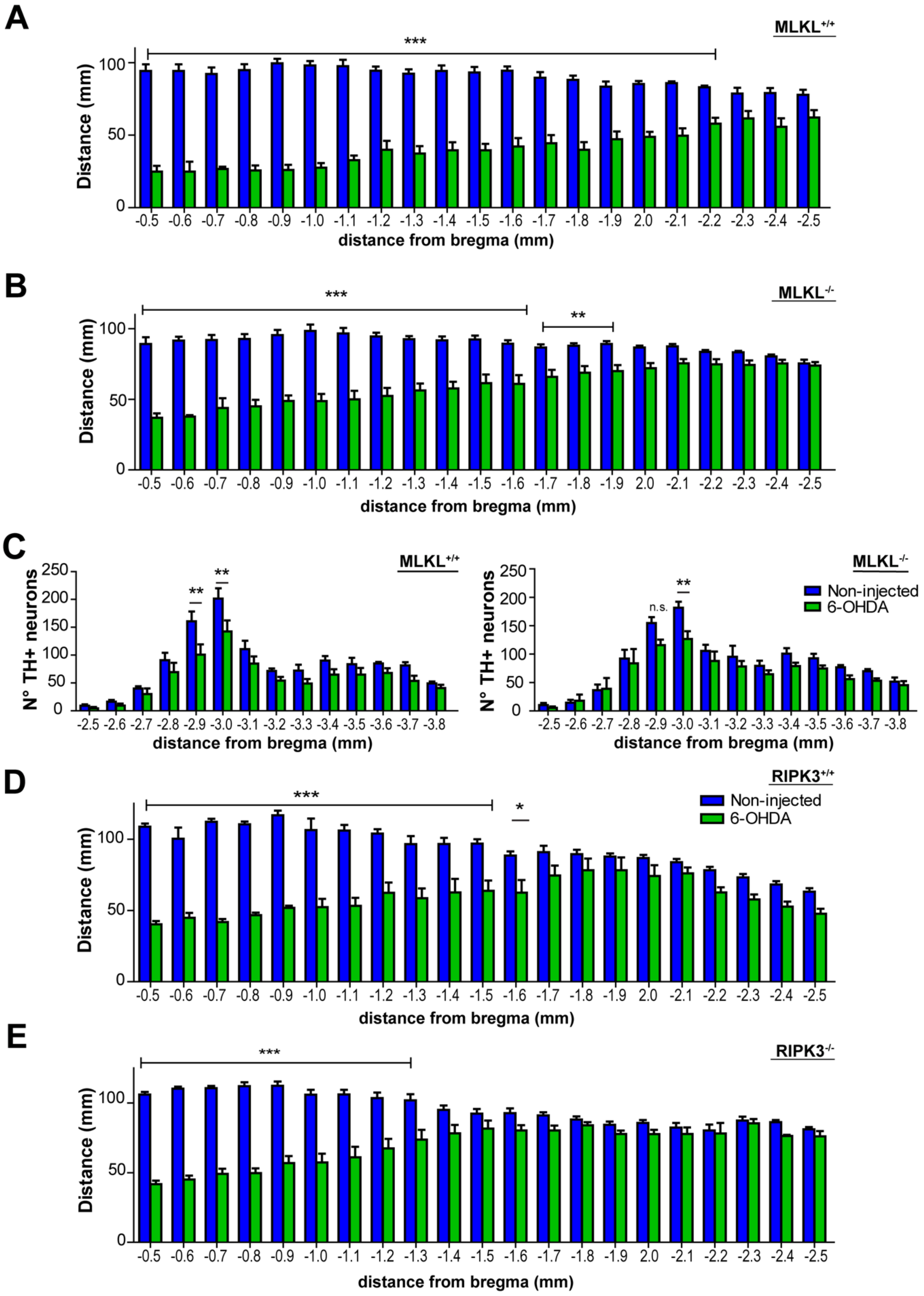
A
MLKL+/+
***
Distance (mm)
100
50
0
-0.5 -0.6 -0.7 -0.8 -0.9 -1.0 -1.1 -1.2 -1.3 -1.4 -1.5 -1.6 -1.7 -1.8 -1.9 2.0 -2.1 -2.2 -2.3 -2.4 -2.5
distance from bregma (mm)
B
MLKL-/-
***
**
Distance (mm)
100
50
0
-0.5 -0.6 -0.7 -0.8 -0.9 -1.0 -1.1 -1.2 -1.3 -1.4 -1.5 -1.6 -1.7 -1.8 -1.9 2.0 -2.1 -2.2 -2.3 -2.4 -2.5
distance from bregma (mm)
C
MLKL+/+
**
**
N° TH+ neurons
250
200
150
100
50
0
-2.5 -2.6 -2.7 -2.8 -2.9 -3.0 -3.1 -3.2 -3.3 -3.4 -3.5 -3.6 -3.7 -3.8
distance from bregma (mm)
MLKL-/-
n.s.
**
Non-injected
6-OHDA
N° TH+ neurons
250
200
150
100
50
0
-2.5 -2.6 -2.7 -2.8 -2.9 -3.0 -3.1 -3.2 -3.3 -3.4 -3.5 -3.6 -3.7 -3.8
distance from bregma (mm)
D
RIPK3+/+
***
*
Non-injected
6-OHDA
Distance (mm)
100
50
0
-0.5 -0.6 -0.7 -0.8 -0.9 -1.0 -1.1 -1.2 -1.3 -1.4 -1.5 -1.6 -1.7 -1.8 -1.9 2.0 -2.1 -2.2 -2.3 -2.4 -2.5
distance from bregma (mm)
E
RIPK3-/-
***
Distance (mm)
100
50
0
-0.5 -0.6 -0.7 -0.8 -0.9 -1.0 -1.1 -1.2 -1.3 -1.4 -1.5 -1.6 -1.7 -1.8 -1.9 2.0 -2.1 -2.2 -2.3 -2.4 -2.5
distance from bregma (mm)

**Supplementary Figure 3. MLKL and RIPK3 deficiency delays 6-OHDA-induced axonal neurodegeneration *in vivo* (related to Figure 5). (A-C)** $MLKL^{+/+}$, $MLKL^{-/-}$ and **(D, E)** $RIPK3^{+/+}$, $RIPK3^{-/-}$ mice were injected with 6-OHDA in the right striatum. Left hemisphere was used as non-injected control side. **(A, B)** Axonal lengths were analyzed in serial coronal sections immunostained for TH every 100 μm at 7 days post 6-OHDA injection. Antero-posterior coordinates relative from bregma are shown in the Y axis in $MLKL^{+/+}$, $MLKL^{-/-}$ mice, respectively. **(C)** Spatial distribution of the number of TH-positive neurons in the SNpc in the non-injected and injected hemisphere of $MLKL^{+/+}$ (left graph) and $MLKL^{-/-}$ (right graph) at 7 days post 6-OHDA injection. **(D, E)** Axonal lengths were analyzed in serial coronal sections at 7 days post 6-OHDA injection in $RIPK3^{+/+}$, $RIPK3^{-/-}$ mice, respectively. Data are shown as mean ± SEM. Statistical differences were obtained using two-way ANOVA, followed by Bonferroni's *post-hoc* test. * $p < 0.05$, ** $p < 0.01$; *** $p < 0.001$. n.s= non-significant. n = 8 animals per group.

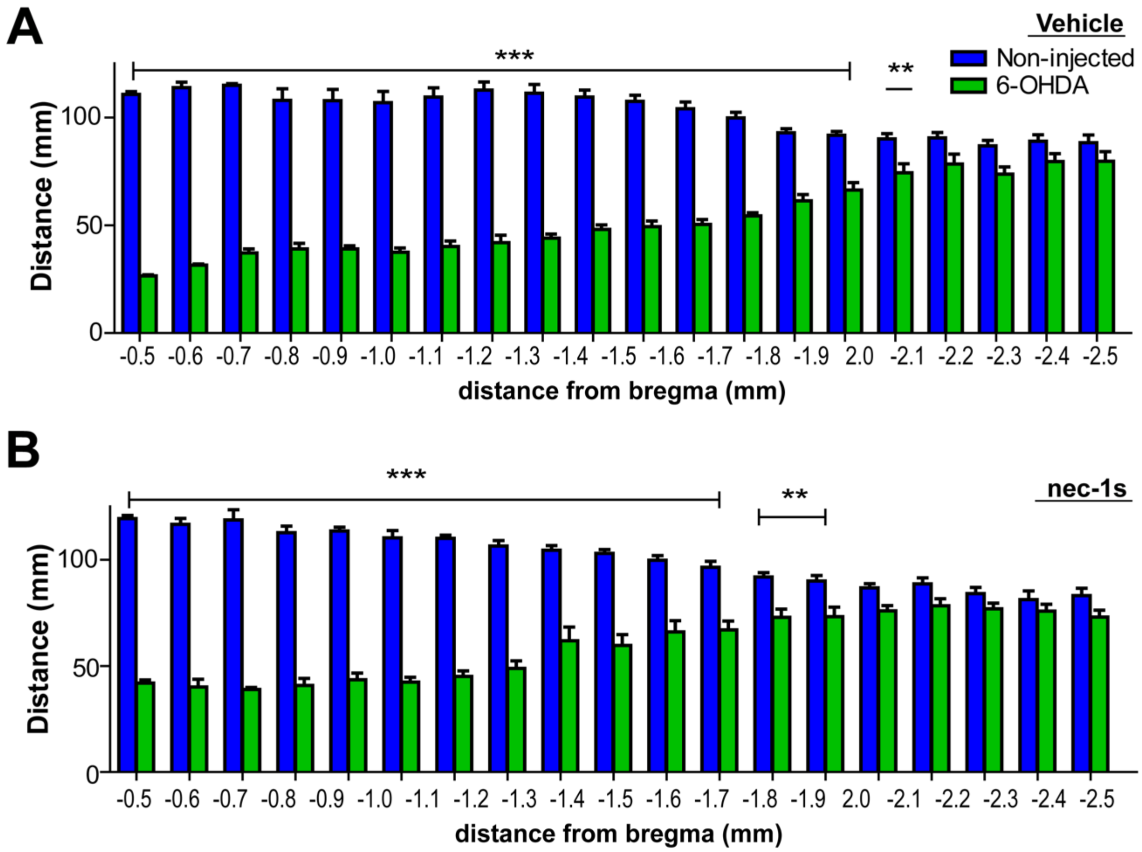


**Supplementary Figure 4. Pharmacological inhibition of RIPK1 delays axonal degeneration after 6-OHDA injection (related to Figure 7).** WT mice were pre-treated with nec-1s for 3 days and the unilaterally injected with 6-OHDA in the right striatum. Then the mice were followed by 7 days with daily injections of nec-1s. Vehicle (Veh) treatment was used as a control for the nec-1s injections. Axonal lengths were analyzed in serial coronal sections immunostained for TH every 100 μm at 7 days post 6-OHDA injection in Veh **(A)** or nec-1s **(B)** treated mice. Antero-posterior coordinates relative from bregma are shown in the Y axis. Data are shown as mean ± SEM. Statistical differences were obtained using two-way ANOVA, followed by Bonferroni's *post-hoc* test. ** $p < 0.01$; *** $p < 0.001$. n = 9 animals per group.
